# Bordetella pertussis BctCBA Mediates Citrate-Dependent Zn^2+^ and Ni^2+^ binding

**DOI:** 10.64898/2026.07.27.740893

**Authors:** Lívia Barreiro Chiorato, Mariana Silveira Derami, João Pedro Aroucha, Lucas Rodrigo de Souza, Natalia Fernanda Bueno, Katlin Brauer Massirer, Mario Henrique Bengtson, Germán Gustavo Sgro, Marilis do Valle Marques, Rafael Junqueira Borges, Leonardo Talachia Rosa

**Affiliations:** Departamento de Bioquímica e Biologia Tecidual. Laboratório de Bioquímica de Complexos Bacterianos. Instituto de Biologia. Universidade Estadual de Campinas (UNICAMP) Monteiro Lobato, 255 Campinas 13083-862 Brasil; Center for Medicinal Chemistry - CQMED, Structural Genomics Consortium (SGC), UNICAMP, Campinas,13083-886 Brazil; Center for Molecular Biology and Genetic Engineering - CBMEG, UNICAMP, Campinas, 13083-875 Brazil; Departamento de Ciências Biomoleculares, Faculdade de Ciências Farmacêuticas de Ribeirão Preto, Universidade de São Paulo, Ribeirão Preto, 14040-903 Brazil; Departamento de Microbiologia, Instituto de Ciências Biomédicas, Universidade de São Paulo, São Paulo, 05508-900 Brazil

## Abstract

*Bordetella pertussis*, the causative agent of whooping cough, is a reemerging public health threat. While the Tripartite Tricarboxylate Transporter (TTT) system BctCBA was previously implicated solely in citrate uptake, we demonstrate that the solute-binding protein BctC specifically binds citrate chelated with Zn²⁺ and Ni²⁺. To elucidate the molecular mechanism of this interaction, we determined the crystal structures of BctC in three states: apo, open, and closed (citrate–zinc-bound), defining the structural determinants for metal-citrate recognition. Comparative analyses suggest that citrate-mediated divalent cation binding is a widespread feature among bacterial TTT homologs. Finally, *in silico* modeling of the full BctCBA complex predicts an elevator-type transport mechanism. Together, these findings redefine the functional scope of BctCBA, revealing a sophisticated strategy by which *B. pertussis* exploits organic chelators to acquire essential trace metals during infection.

## Introduction

Pertussis, or whooping cough, is a highly contagious respiratory disease caused by the Gram-negative bacterium *Bordetella pertussis*. It primarily affects children under the age of four and is characterized by persistent coughing fits, which can lead to lung tissue damage and necrosis. Despite the availability of vaccines, pertussis outbreaks have re-emerged over the past decade, prompting the classification of *B. pertussis* as a re-emerging pathogen (Melvin *et al*, 2014).

The genome of *Bordetella* species encodes an overrepresented family of uptake systems known as Tripartite Tricarboxylate Transporters (TTTs) (Antoine *et al*, 2003). TTTs are one of three major families of high-affinity, solute-binding protein (SBP)-dependent transporters in bacteria, alongside ATP-Binding Cassette (ABC) and Tripartite ATP-independent Periplasmic (TRAP) transporters. These systems function through periplasmic SBPs that bind target ligands with high affinity, followed by delivery to membrane transport complexes. The transport process is powered either by ATP (ABC) or an electrochemical gradient (TRAP and TTT) (Rosa *et al*, 2018)

TTT systems are composed of three components: TctA, a 12-transmembrane-domain transporter; TctB, a four-transmembrane protein of unknown function; and TctC, a periplasmic SBP with high ligand specificity. A detailed review of the TTT systems is available (Rosa *et al*, 2018). Remarkably, the genome of *B. pertussis* Tohama I encodes 78 *tctC* homologs, making it the most abundant gene family in the genome. In contrast, only one complete operon is observed, the *Bordetella* citrate transporter operon *bctCBA* (Antoine *et al*, 2003).

Citrate was the first ligand identified for a TTT system, originally in the *tctCBA* operon of *Salmonella typhimurium* (Sweet *et al*, 1979), and remains the most commonly reported ligand among TTT systems (Rosa *et al*, 2018). In *B. pertussis*, citrate induces expression of the *bctCBA* operon, and mutations in these genes abolish citrate uptake (Antoine *et al*, 2003, 2005). The SBP BctC interacts with both BctA, to mediate citrate transport, and the sensor protein BctE, part of the constitutively expressed two-component system *bctDE*, which regulates *bctCBA* expression (Antoine *et al*, 2005).

In this study, we provide evidence that the BctCBA system functions as a high-affinity transporter of divalent cations in association with citrate. We demonstrate that the divalent cations Zn²⁺ and Ni²⁺ enhance the thermal stability of BctC and increase its affinity for citrate. Through the determination of crystallographic structures in three distinct states, we show that the transition from the open to the closed conformation is triggered by the presence of zinc and does not occur in the presence of citrate alone. Furthermore, we present evidence suggesting that the exploitation of citrate’s metal-chelating properties for the high-affinity uptake of divalent cations may represent a widespread functional strategy among bacterial members of the TTT transporter family. Finally, we use *in silico* modeling to propose an elevator-type transport mechanism for the TTT family.

## Results

### BctC affinity for citrate is enhanced in the presence of the divalent cations Zn^2+^ and Ni^2+^

We set out to investigate the *in vitro* ligand-binding properties of BctC (Fig 1). As a periplasmic protein, BctC is partially exposed to environmental pH variations. Therefore, we first used differential scanning fluorimetry (DSF) to determine the optimal pH for its stability, selecting pH 7.5 for subsequent analyses (Fig. EV1A). BctC has previously been characterized as a citrate-binding protein; because citrate is a highly abundant intracellular metabolite and may have co-purified with BctC, to ensure removal of any bound ligand we denatured the protein via dialysis in 6 M urea, and subsequently refolded it. Circular dichroism analysis confirmed that the secondary structure of BctC remained preserved after this refolding process (Fig. EV1B). To explore potential additional ligands, we performed a DSF-based screening of 85 metabolites (including amino acids, vitamins, and carboxylic acids) monitoring for thermal stabilization effects (Appendix Table 1). Among all tested compounds, only citrate induced thermal stabilization of BctC above the threshold (ΔTm ≥ 2 °C). Finally, size-exclusion chromatography revealed that BctC elutes as a monomer in both the presence and absence of citrate (Fig. 1B).

**Figure 1:**
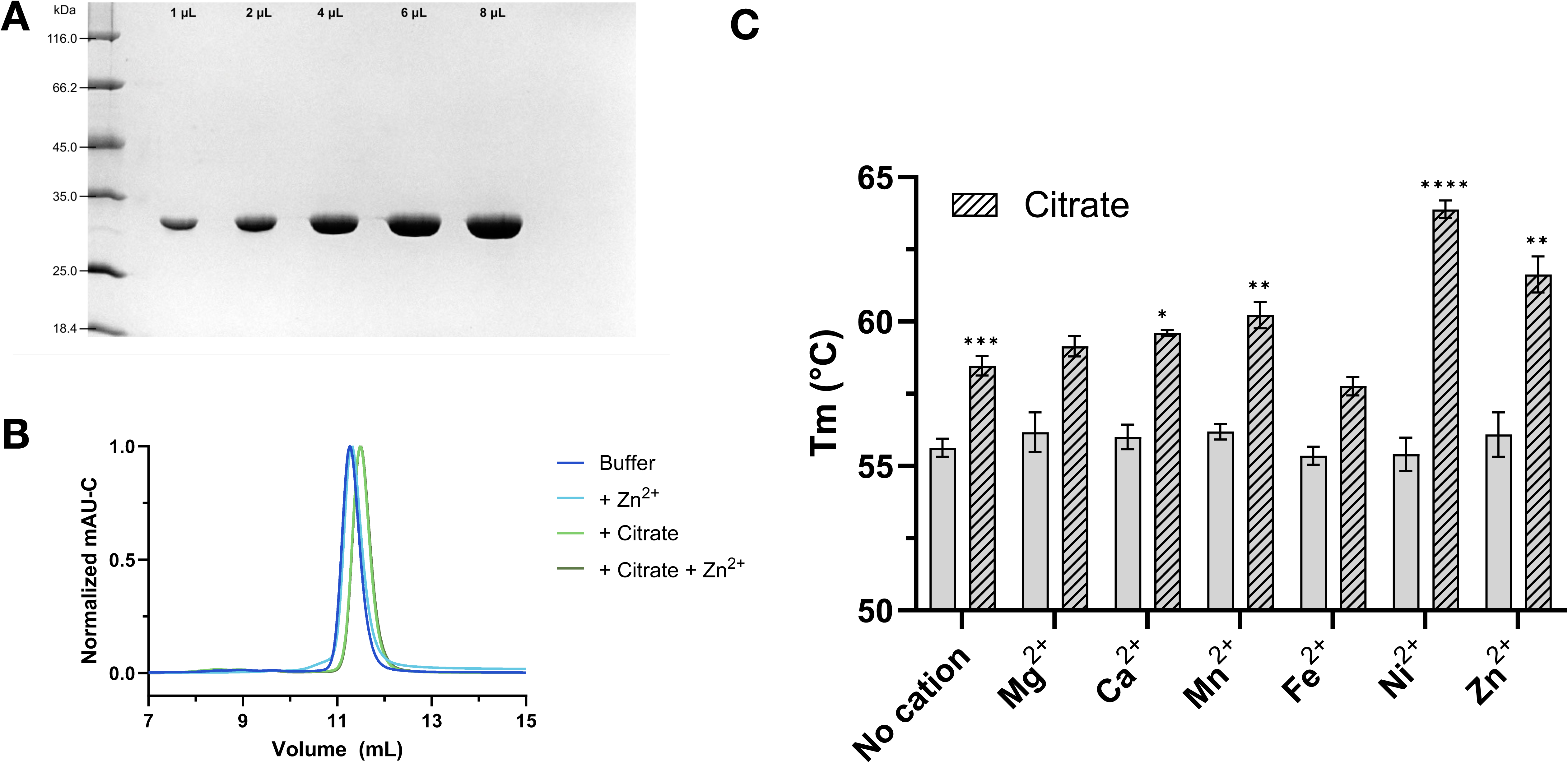
Biochemical characterization of BctC and its interactions with citrate and divalent cations. (a) Increasing amounts of purified BctC analyzed by 12% SDS–PAGE following urea dialysis and TEV protease cleavage to assess protein purity. (b) Size-exclusion chromatography profiles of BctC on a Superdex 200 Increase 16/300 column. Equal amounts of protein were injected in the buffer alone, with zinc chloride, sodium citrate, or sodium citrate plus zinc chloride. Elution volumes (estimated molecular masses) were 11.3 mL (33.3 kDa) for buffer alone, 11.3 mL (32.8 kDa) for zinc chloride, 11.5 mL (30.7 kDa) for sodium citrate, and 11.5 mL (30.7 kDa) for sodium citrate plus zinc chloride. (c) Differential scanning fluorimetry (DSF) analysis of BctC in the presence of divalent cations alone or citrate supplemented with divalent cations. Experiments were performed in triplicate. Statistical analyses were performed using Welch’s t-test (two-tailed), assuming unequal variances between groups. Each cation condition was compared with the buffer control, and each citrate + cation conditions were compared with citrate control. Asterisks indicate statistically significant differences (*p< 0.05; ** p<0.01; *** p<0.001; ****p<0.0001).

Citrate has previously been described as a chelating agent for divalent cations (Krom *et al*, 2000; Banerjee *et al*, 2016). Based on this, we hypothesized that the BctCBA system may function not only in the uptake of citrate as a carbon source but also in scavenging biologically important divalent cations, particularly in the context of bacterial pathogenesis. To test this, we conducted a new round of DSF experiments, adding divalent cations at a concentration of 10 mM. The addition of Ca²⁺, Mn²⁺, Ni²⁺ and Zn²⁺ led to a statistically significant increase in BctC melting temperature, compared to citrate alone, whereas Fe²⁺ and Mg²⁺ did not produce a significant effect (Fig. 1C).

To investigate the impact of the divalent ion binding, we measured the BctC affinity for citrate in the presence of the various divalent cations using isothermal titration calorimetry (ITC) . Consistent with the DSF results, the presence of Ni²⁺ and Zn²⁺ increased BctC’s affinity for citrate, with the dissociation constant (*K*_d_) decreasing from 4.23 µM to 0.88 µM and 2.04 µM, respectively. In contrast, the presence of Fe²⁺, Ca²⁺, Mg²⁺, and Mn²⁺ resulted in reduced affinity. Although a decrease in affinity might seem unexpected, it is likely explained by citrate’s chelation of these metal ions, which reduces the availability of free citrate and leads to an apparently higher K_d_ (Table 1, Fig. EV2 and Appendix Table 2).

**Table 1:** Isothermal Titration Calorimetry plots of BctC in the presence of citrate. Corrected heat rates are shown in the top panel, and normalised fits are shown in the bottom. Divalent ions were included in the dialysis and assay buffers and were present at identical concentrations in both the ITC cell and syringe.

| *Ligand\*\** | *K<sub>d</sub>* (μM) | $\Delta H$<br>(kcal mol<sup>-1</sup>) | $-T\Delta S$<br>(kcal mol<sup>-1</sup>) | $\Delta G$<br>(kcal mol<sup>-1</sup>) | *N* |
| --- | --- | --- | --- | --- | --- |
| Citrate | 4.23 ± 0,43 | 13.57 ± 0.11 | -20.90 | -7.34 | 0.95 ± 0.03 |
| * Citrate + Mg<sup>2+</sup> | 24.95 ± 6,16 | 11.55 ± 1.38 | -17.85 | -6.29 | 0.84 ± 0.08 |
| Citrate + Ca<sup>2+</sup> | 27.00 ± 3,69 | 16.50 ± 0.82 | -22.77 | -6.24 | 0.73 ± 0.02 |
| Citrate + Mn<sup>2+</sup> | 16.07 ± 1,27 | 11.73 ± 0.48 | -18.30 | -6.55 | 0.73 ± 0.02 |
| * Citrate + Fe<sup>2+</sup> | 17.65 ± 1,69 | 14.65 ± 0.38 | -21.15 | -6.49 | 0.68 ± 0.01 |
| Citrate + Ni<sup>2+</sup> | 0.88 ± 0,25 | 8.14 ± 0.36 | -16.43 | -8.28 | 0.72 ± 0.01 |
| Citrate + Zn<sup>2+</sup> | 2.04 ± 0,46 | 6.67 ± 0.29 | -14.43 | -7.80 | 0.74 ± 0.01 |

\* For these conditions, statistical analysis was performed using biological duplicates instead of triplicates.

### Crystal structures of BctC reveal Zn²⁺-dependent stabilization of the closed conformation

To investigate the binding mechanism of BctC, crystal structures were determined in the absence of ligand (apo form), as well as in the presence of citrate alone, or citrate and zinc chloride (Table 2). The apo structures and the structure crystallized with citrate alone both adopt an open conformation. In the latter, electron density indicative of partial citrate occupancy is observed within domain 1 of the binding pocket, suggesting that this binding is only weakly stabilized (Fig. EV3A). In contrast, in the presence of both citrate and zinc, BctC adopts a closed conformation, indicating that zinc stabilizes this state in a manner that citrate alone is unable to achieve. Regarding asymmetric unit (ASU) composition, both the apo and open/citrate-bound structures contain a single copy per ASU, whereas the closed BctC/citrate/Zn^2+^ structure forms a tetramer with an RMSD below 0.45 Å between the monomers across the 302 considered atoms. A second apo structure yielded 16 copies per ASU, with RMSD values ranging between 0.35 Å and 0.71 Å across 302 atoms.

**Table 2:**
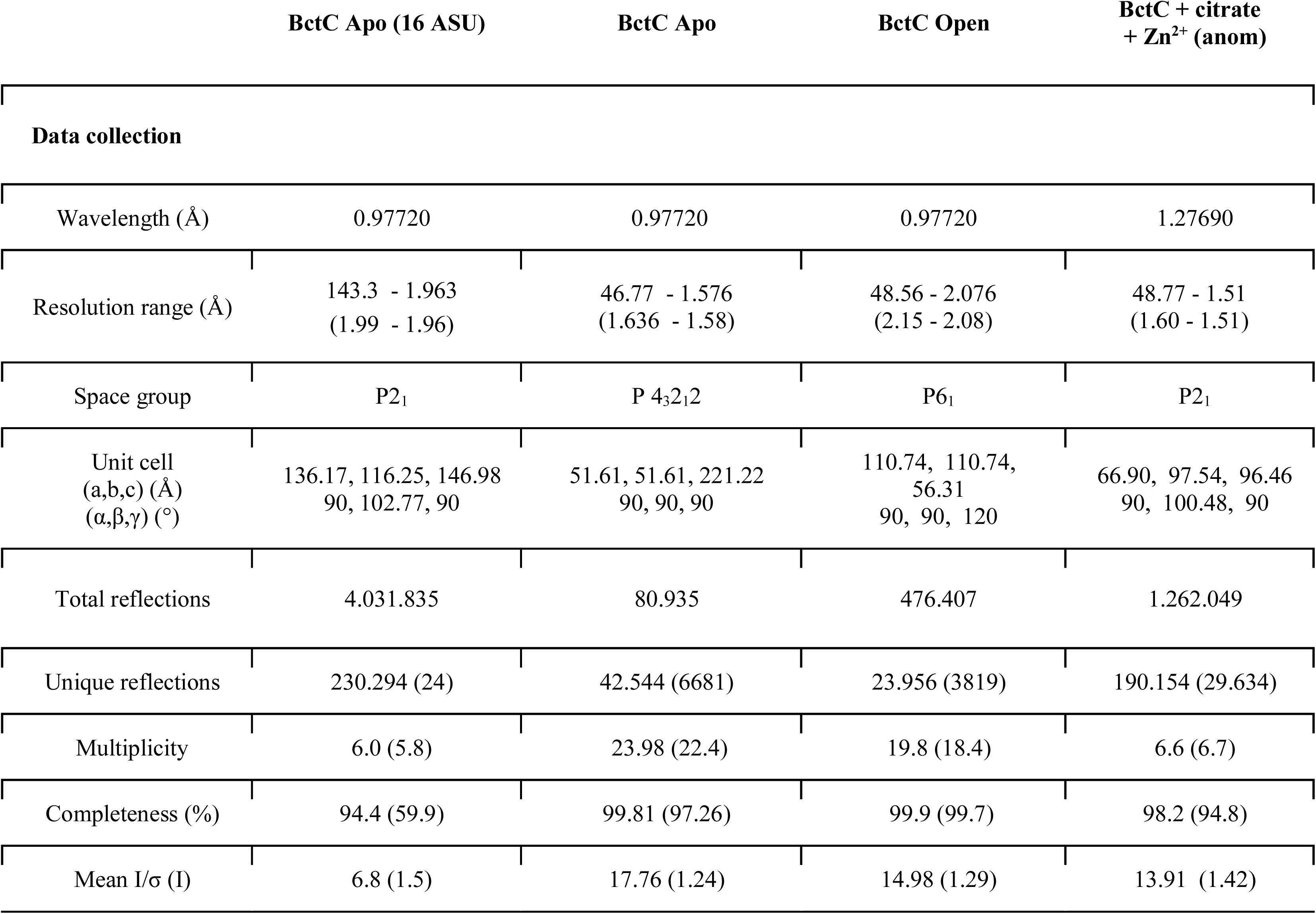
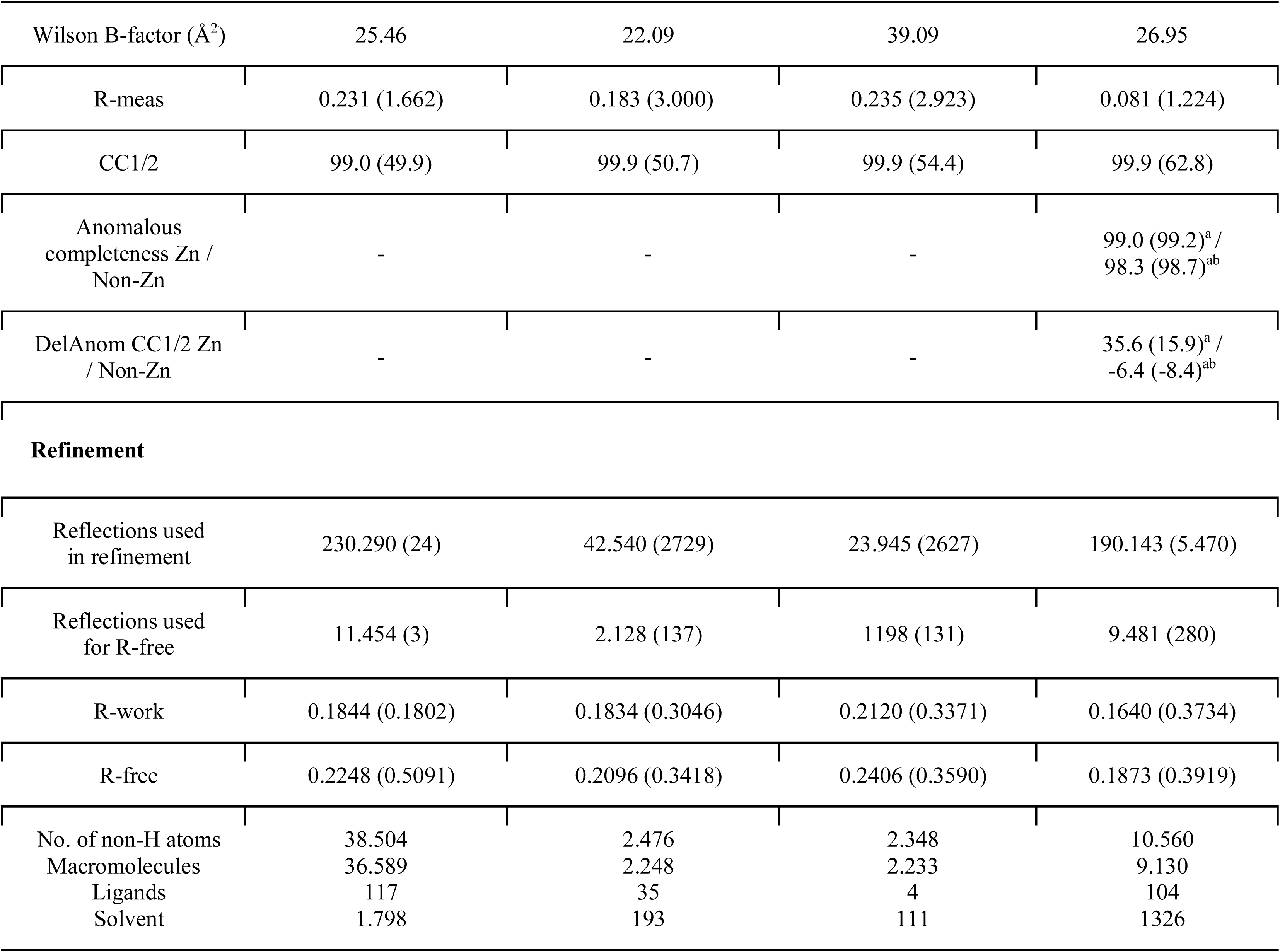
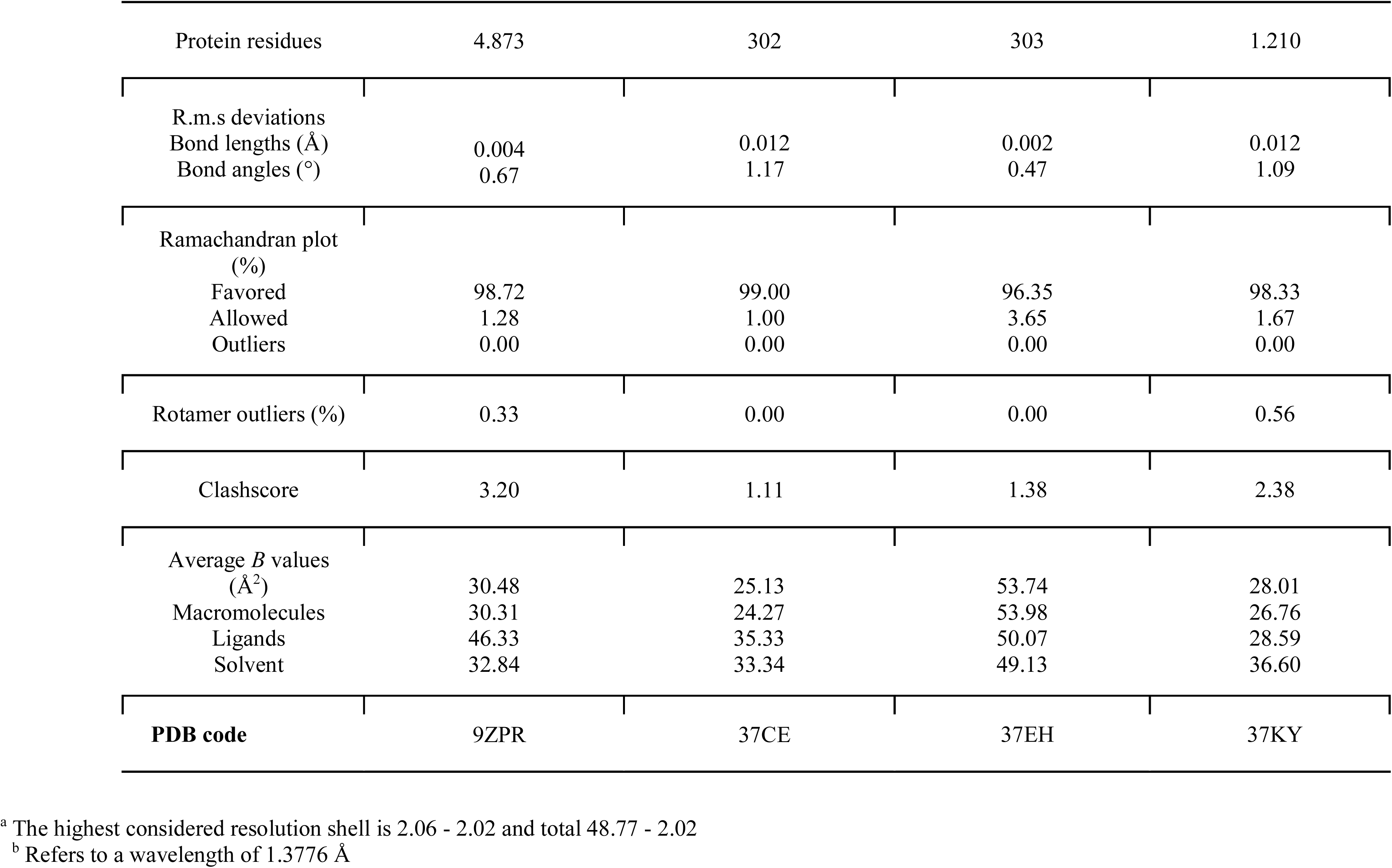
Protein crystallography data collection and refinement statistics for BctC.

BctC is organized into two globular domains connected by a hinge and separated by a binding cleft, forming a Venus flytrap–like architecture (Fig. 2A). It is classified as a subclass E2 soluble binding protein (SBP), together with all TTT SBPs characterized to date (Scheepers *et al*, 2016). The two domains adopt the canonical fold observed in the TTT family. Domain 1 comprises residues 31–131 and 260–334 and consists of a five-stranded β-sheet (β2–β1–β3–β9–β4) surrounded by six α-helices and one short helix. Cysteines C38 (β1) and C52 (α1) are located in close proximity. In all models, they appear partially reduced, with double occupancy, likely due to cytoplasmic expression and the reducing conditions used during purification. Domain 2 comprises residues 132–259 and contains a hydrophobic core formed by five β-strands (β6–β5–β7–β4–β8), surrounded by four α-helices and three short helices (Fig. 2B).

**Figure 2:**
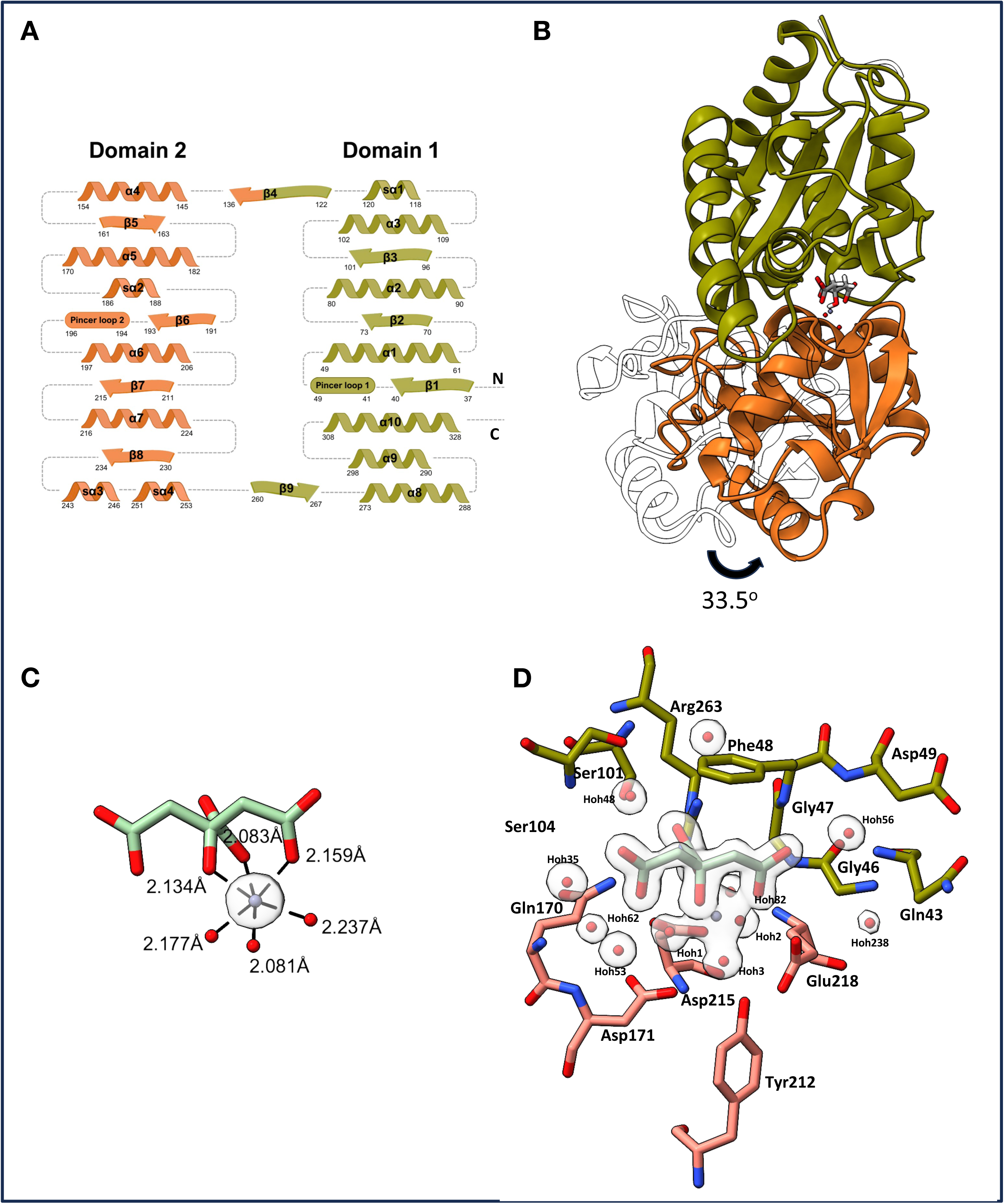
Structural basis of citrate and zinc recognition by BctC. (a) Domain organization and secondary-structure architecture of BctC. α-helices are represented as spirals, β-strands as arrows, and the pincer loops involved in substrate coordination as cylinders. Residue numbers are indicated at the beginning and end of each secondary-structure element. Domain 1 is shown in olive green and Domain 2 in orange. The N- and C-termini are labeled. (b) Structural comparison between the apo and holo crystal structures of BctC. Both structures are aligned using Domain 1, shown in olive green. Domain 2 is shown in orange for the holo structure and as a white outline for the apo structure, highlighting the 33.5° conformational shift between the two states. Citrate from the holo structure is shown as sticks, with carbon atoms in gray, oxygen atoms in red, and hydrogen atoms in white. Water molecules are represented as red spheres, and the zinc ion as a purple sphere. (c) Zn^2+^ coordination within the binding pocket of BctC. Zn^2+^ is shown in cyan, with oxygens in red. Waters are depicted as red spheres. Distances are depicted as dark dashed lines. The polder electron density map is displayed as a transparent gray density, contoured at 15 RMSD. (d) Three-dimensional view of citrate and zinc coordination in the BctC holo structure. Oxygen atoms are shown in red, nitrogen atoms in blue, water molecules as red spheres, and the zinc ion as a purple sphere. Citrate carbon atoms are colored cyan. Carbon atoms from BctC residues belonging to Domain 1 are shown in olive green, whereas those from Domain 2 are shown in orange. The polder electron density map is displayed as a transparent gray density, contoured at 7.5 RMSD.

A hinge motion of 33.5° is observed between the apo and closed conformations, using the Cα atoms of D49 (α1 in domain 1), I262 (β9, inter-domain hinge), and G198 (α6 in domain 2) as reference points (Fig. 2B). Closure around the ligand occurs largely via a rigid-body motion, as indicated by low RMSD values of separated domains between open and closed states (0.6 Å for domain 1 and 1.0 Å for domain 2). Nevertheless, a subtle local rearrangement is observed in the region between β6 and α6, likely in response to coordination of pincer loop 2 by water molecule 238 (see below).

Zinc adopts an octahedral geometry, coordinated primarily by citrate through the carboxyl groups at C6 and C1, and the hydroxyl group at C3 (Fig. 2C). In addition, it is stabilized by three water molecules that form hydrogen bonds with both citrate and domain 2 residues within the binding pocket. Water 1 bridges interactions with Asp215 and Asp171. Water 2 bridges interactions with Asp215 and Glu218. Water 3 bridges interactions with Asp171, with the backbone of Gly197 and with Tyr212, which projects its side chain toward the binding pocket (Fig. 2D).

The assignment of zinc ions was further supported by anomalous difference peaks observed in a dataset collected near the Zn absorption edge and by their marked reduction or absence in a dataset collected below the edge (Table 3). This analysis also revealed an additional zinc ion mediating interactions among three different chains within the asymmetric unit (ASU). This ion coordinates His90 from chains C and D and Glu31 from chain A, with a fourth coordination site occupied by a water molecule, completing a tetrahedral geometry. Size-exclusion chromatography of BctC in the presence of citrate and zinc shows that the protein continues to elute as a monomer (Fig 1B), suggesting that this inter-chain zinc coordination is likely a crystallographic artifact without biological relevance.

**Table 3:**
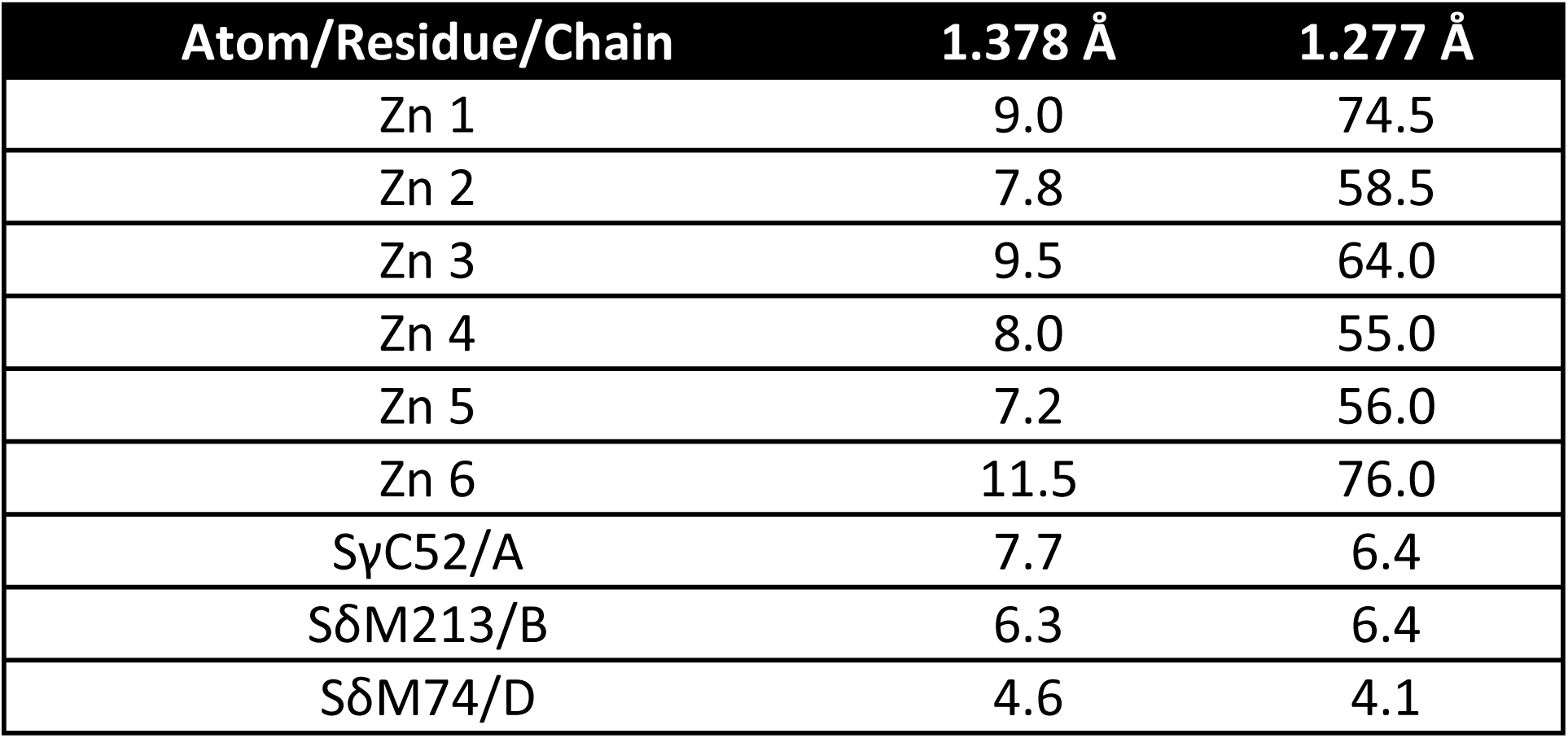
Anomalous peak heights (RMSD) from anomalous difference Fourier maps of BctC/citrate/Zn^2+^ crystallographic structure. Peaks above the 2.5 RMSD threshold are shown.

Coordination of citrate within the binding pocket is described as follows (Fig. 2D). As conserved among TTT family SBPs, Phe48 serves as a docking site for the hydrophobic backbone of the ligand, contacting citrate carbons C2 and C4, together with Ile78. The carboxyl group at C1 forms hydrogen bonds with the backbone of Gly197, part of pincer loop 2, and with Phe48, in addition to water-mediated interactions involving waters 2 and 3. Water 56 provides further stabilization by bridging hydrogen bonds with Gln43, Gly46, and Asp49. This water molecule is one of two conserved waters in TTT SBPs that mediate interactions between ligand carboxyl groups and the flexible loops at the entrance of the binding pocket, which together form a “pincer-like” structure (P41–D50 in domain 1 and F194–G197 in domain 2), thereby structuring these loops (Rosa *et al*, 2018). Notably, this water molecule is also present in the apo structure, suggesting a role in loop pre-organization and initial ligand recognition. The second conserved water molecule involved in this coordination (water 238) is present only in the closed structure and interacts exclusively with the protein, bridging the two domains, rather than with the ligand. This is likely due to displacement of the ligand carboxyl group relative to other TTT homologs, toward zinc coordination. This water is absent in the apo structure, indicating its importance in stabilizing domain 2 following ligand recognition.

The carboxyl group at C5 is coordinated primarily via hydrogen bonds with Gln170 and water-mediated interactions involving waters 35, 48, and 62. Water 35 forms hydrogen bonds with Asn107 and the backbone of Gly103 and Gln170. Water 48 bridges interactions with Asn107 and the backbones of Ser104 and Ile78. Water 62 bridges hydrogen bonds with the backbone of Gly164 and Asp171. The carboxyl group at C6 is coordinated via a salt bridge with Arg263 and hydrogen bonds with Ser101, in addition to interactions with waters 1, 2, and 3. The hydroxyl group at C3 participates primarily in zinc coordination and water-mediated interactions.

Finally, all six α-helices surrounding the binding pocket are oriented with their N-termini facing the pocket, contributing to stabilization of the negatively charged ligand via the helix dipole effect (Fig. EV3B).

In summary, the crystallographic structures of BctC in three different states suggest that citrate alone is insufficient to trigger domain closure in BctC, leaving the protein in an open conformation with only partial ligand occupancy. Following ligand recognition in domain 1, complete domain closure and full stabilization of the binding pocket require the synergistic coordination of zinc, which bridges citrate with domain 2 residues and induces the necessary hinge motion.

### Comparative Analysis of Citrate and Divalent Cation Binding Across BctC Homologs

Citrate has been recognized as the predominant substrate of Tripartite Tricarboxylate Transporters (TTT) (Rosa *et al*, 2018). To investigate whether the uptake of divalent cations is a conserved feature across this transporter family, we cloned and expressed BctC homologs from five organisms for which experimental evidence of citrate binding or transport has been reported: *Salmonella enterica* serovar Typhimurium (_SE_TctC) (Sweet *et al*, 1979), *Corynebacterium glutamicum* (_CG_TctC) (Brocker *et al*, 2009), *Geobacillus thermodenitrificans* (_GT_TctC) (Graf *et al*, 2016), *Advenella* sp. (_AV_TctC) (Schäfer *et al*, 2019), and *Pseudomonas aeruginosa* (_PA_TctC) (Underhill & Cabeen, 2022) (Fig. 3A).

**Figure 3:**
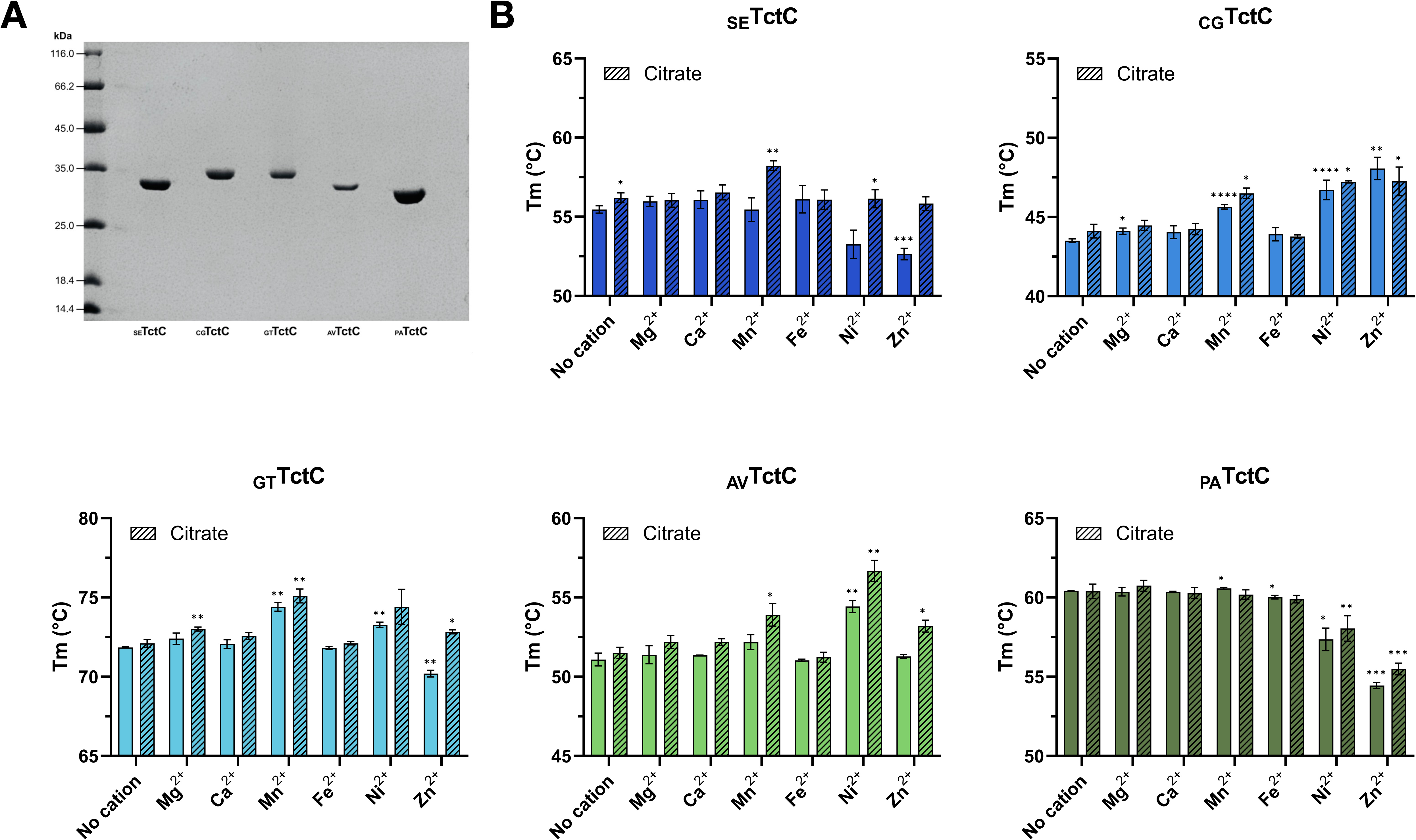
Analysis of five citrate binding TctC homologs across bacterial species. (a) 12% SDS-PAGE gel showing purification of each homolog, after affinity chromatography, urea denaturation and renaturation and size-exclusion chromatography. (b) Differential scanning fluorimetry (DSF) analysis of TctC homologs in the presence of divalent cations alone or citrate supplemented with divalent cations. Experiments were performed in triplicate. Statistical analyses were performed using Welch’s t-test (two-tailed), assuming unequal variances between groups. Each cation condition was compared with the buffer control, and each citrate + cation conditions were compared with citrate control. Asterisks indicate statistically significant differences (*p< 0.05; ** p<0.01; *** p<0.001; ****p<0.0001).

Differential scanning fluorimetry (DSF) analyses revealed that _SE_TctC undergoes a significant increase in melting temperature in the presence of both citrate and Mn²⁺, whereas citrate or Mn²⁺ alone produced little or no effect. Similarly, _AV_TctC exhibited significant thermal stabilization when citrate was combined with Zn²⁺, Mn²⁺, or Ni²⁺ compared with either citrate or the corresponding metal ion alone (Fig 3B). These two homologs share the highest sequence identity with BctC, with 64% and 83.5%, respectively, while _PA_TctC, _GT_TctC and _CG_TctC share 60.4%, 32.3% and 28.5%, respectively, suggesting a common ancestrality among those showing ion-dependent thermal stabilization. Residues directly involved in Zn^2+^ coordination in BctC are conserved among the homologs, with the exception of _CG_TctC.

Notably, several of the TctC homologs tested displayed either thermal stabilization or destabilization in the presence of divalent cations alone. These effects likely result from direct interactions between the ions and the proteins, independent of citrate binding. Overall, these results suggest that synergistic recognition of citrate and divalent cations may be conserved in at least a subset of TTT systems, although the extent and specificity of this phenomenon vary among homologs.

### In silico evidence for an elevator-type mechanism in the TTT family

The mechanism by which members of the Tripartite Tricarboxylate Transporter (TTT) family translocate ligands across the cytoplasmic membrane remains elusive. Here, we use the BctCBA complex as a model system to investigate this process. Structural models of BctCBA were generated *in silico* using the AlphaFold3 server (Abramson *et al*, 2024), with two Na⁺ ions and one citrate molecule included as inputs. Five models were predicted with high confidence, with an interface predicted TM-score (ipTM) of 0.85, a predicted TM-score (pTM) of 0.86, and most regions of the complex displaying high confidence (pLDDT > 90). Predicted aligned error (PAE) plots further support the overall domain organization and inter-subunit arrangement (Fig EV4A). Since BctC is a periplasmic binding protein and was invariably modelled in the same position relative to BctA, we will refer to the BctC-bound side of BctA as the periplasmic side, and the opposite side as the cytoplasmic side.

Looking from the cytoplasmic to the periplasmic side of the models, two asymmetric domains are distinguishable, separated by a central cleft (Fig. 4A). This topology is suggestive of an elevator-type translocation mechanism, characteristic of the Ion-transport Superfamily (ITS), in which one domain alternates the exposure of a ligand binding site between the periplasm and cytoplasm (elevator domain), while the second domain shows little conformational change (scaffold domain). Example of bacterial transport families operating via such mechanism are the Divalent anion Sodium Symporter (DASS/SLC13) and the ATP-Independent Periplasmic (TRAP) transporters (Davies *et al*, 2024; Sauer *et al*, 2020). Domain 1, the elevator domain, is composed by BctA residues 1-144 and 233-392 (Fig. 4A and 4B, cyan), while domain 2, the scaffold domain, is formed by residues 145-231 and 393-507 (Fig. 4A and 4B, green). BctB is modelled adjacent to domain 2, and might serve as an extension of the scaffold domain (Fig. 4A and 4B-blue), stabilizing it, analogous to what is described in TRAP transporters (Peter *et al*, 2022). BctA models were obtained in two different conformational states, with cytoplasm facing (models 0 to 3)(Fig. 4B, left panel) and periplasm facing (model 4) surfaces (Fig. 4B, right panel). For clarity, these models will be referred to as _in_BctBA and _out_BctBA, respectively. BctC was invariably modelled in the open conformation, and the citrate molecule was either bound to BctA (model 0) or BctC (models 1 to 4).

**Figure 4:**
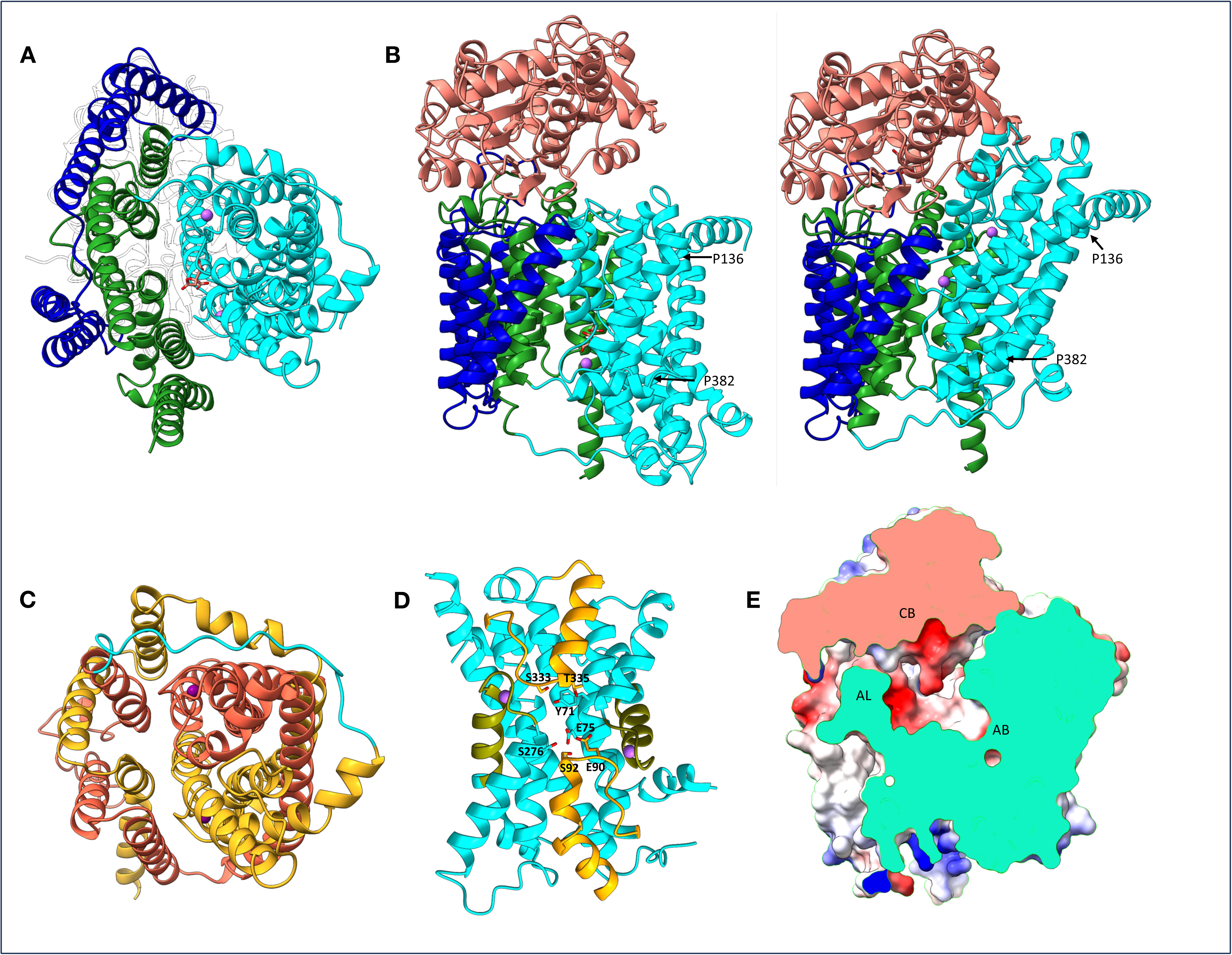
Structural analysis of *in silico* BctCBA complex models. (a) Orthogonal views of _in_BctCBA complex, illustrating the organization of the membrane components into two domains separated by a central cleft. The putative BctA elevator domain is shown in cyan, the BctA scaffold domain in green, and BctB in blue. BctC is shown as a white surface outline in the background. Sodium ions are represented as purple spheres, and citrate is shown in stick representation.(b) BctCBA complex models in two distinct conformational states. _in_BctCBA (left) depicts BctA in a resting, cytoplasm-facing conformation, whereas _out_BctCBA (right) depicts BctA in a periplasm-facing conformation. This conformational transition appears to be enabled by bending or stretching of two α-helices at positions P136 and P382. The color scheme is the same as in panel (a), except that BctC is shown in salmon. (c) Putative pseudo-symmetrical organization of BctA. Orthogonal views of _in_BctA highlight topologically mirrored regions, shown in gold and orange. (d) Putative ligand-coordination motifs in BctA. The elevator domain is shown, with the mirrored hairpins colored orange and the clamshell structures formed by the conserved motif G-Hy₃-∗G-Hy₃-∗G-Hy₂-∗P-G-Hy-G shown in olive green. Residues predicted to contribute to the acidic character of the putative citrate-binding site are shown in stick representation. Sodium ions are depicted as purple spheres. (e) Central slice through _out_BctCBA shown in surface representation, suggesting a potential continuity between the binding pockets of BctC in its open conformation and _out_BctA. Surfaces are colored according to electrostatic potential, with acidic regions shown in red, neutral regions in white, and basic regions in blue. Internal surfaces exposed by the slice are colored cyan for BctA (elevator and scaffold domains) and salmon for BctC. CB, BctC binding pocket; AB, BctA binding pocket; AL, BctA periplasmic loop.

BctA models present a structural pseudosymmetry, encompassing both domains, resulting in two inverted repeats (half 1 - residues L18 to E227 and half 2 - residues L247 to I463) (Fig. 4C). In each, a small helix reaches the middle of the putative transmembrane space, and then turns back to the space of origin, resulting in mirrored hairpins (Hp_in_ - L85 to R105 and Hp_out_ - T327 to I348) (Fig. 4D- orange). Representing the elevator domain in a columbic surface charge potential, a polar, acidic pocket is evidenced among the hydrophobic transmembrane section in between the mirrored hairpins (Fig. EV4B), modelled in model 0 as the citrate binding pocket. Contributing to the negative charge are mainly E75 and Y71, present in helix 4, in addition to E90 and S92 on HP_in_ and S276, S333 and T335 in HP_out_ (Fig. 4D). Adjacent to the inverted hairpins are the two mirrored loops formed by the conserved motifs G-Hy3-∗G-Hy3-∗G-Hy2-∗P-G-Hy-G (G27 to G41 and S260 to G274) (Fig.5D-olive)(Winnen *et al*, 2003). The two predicted Na^+^ ions are bound to a clamshell formed in the interface of these motifs and the hairpins (Fig. 4D-purple). The scaffold domain is not predicted to directly interact with the citrate or sodium binding sites.

In the proposed model, sliding of the elevator domain from _in_BctBA to _out_BctBA occurs via bending of two lateral amphipathic ‘arm’ helices: 50° bending of helix 6b on Proline 136 (close to the periplasmic side) in _in_BctBA and a 41° bending of helix 14c on Proline382 (close to the cytoplasmic side) in _out_BctBA (Fig. 4B). The cytoplasmic loop K228-K248 adjusts its conformation on the other end of the elevator domain to allow for this movement to occur. In _out_BctBA, the putative citrate binding site (Fig. 4E-AB) is not directly exposed to the solvent, but forms a continuous cavity with a long acidic periplasmic loop ( Fig. 4E-AL) of the scaffold domain between helix 8 and 9 (S183 – D208) and the binding pocket of the open conformation of BctC (Fig. 4E-CB).

All models were predicted with BctC on the open conformation. Aligning the crystallographic structures of closed BctC using either the domain1 or domain 2 of the open BctC as reference indicates that the closed conformation could be placed on _in_BctBA, while the positioning of it on _out_BctBA caused large esterical clashes, suggesting the conformational change of BctA could be coupled to the opening of BctC and consequent exposure of its binding pocket (Fig. EV4C).

> Our attempts to model BctCBA in the presence of Zn²⁺ consistently yielded _in_BctBA structures, with Zn²⁺ coordinated by His7 at the N-terminus of BctA. However, the absence of explicitly modelled Zn²⁺ coordination geometry and water molecules—both of which are likely critical for proper citrate binding within the BctA pocket—limits the reliability of these portion of the models. Therefore, we refrain from making further interpretations regarding substrate coordination in BctA.

## Discussion

In this study, we demonstrate that BctC from *B. pertussis*—previously characterized solely as a citrate-binding protein—orchestrates the co-binding of divalent ions alongside citrate. Using recombinant BctC, we confirmed citrate binding via differential scanning fluorimetry (DSF) and measured binding affinities via isothermal titration calorimetry (ITC) in the presence of various divalent cations. Both thermal stabilization and dissociation constants (*K*_d_) indicated statistically tighter citrate binding in the presence of Zn^2+^ and Ni^2+^, but not Fe²⁺ or Mg²⁺. While Ca²⁺ and Mn²⁺ induced thermal stabilization, they did not yield a parallel decrease in *K*_d_ values.

Upon infection, host defenses rapidly deploy mechanisms to deplete blood metal availability—a process termed nutritional immunity. Consequently, pathogens have evolved various nutrient-scavenging mechanisms, essential for successful colonization and infection (Hood & Skaar, 2012; Cerasi *et al*, 2013). Behind iron, zinc is the second most abundant transition metal in cellular metabolism, participating in DNA repair, gene expression, stress response, and virulence. Acting as a potent Lewis acid catalyst, zinc stabilizes reaction intermediates (Andreini & Bertini, 2012), a property crucial for enzymes like the metalloproteinases used to degrade host epithelial barriers (Ramchunder *et al*, 2026). Zinc is also required for proper protein folding, particularly in DNA-binding motifs like the zinc fingers of transcription factors—such as the prototype regulator Ros found in *Agrobacterium tumefaciens* and other Alphaproteobacteria (D’Abrosca *et al*, 2020). Furthermore, zinc buffers against immune-mediated reactive oxygen species and promotes biofilm formation (Wang *et al*, 2021). While zinc metabolism is well documented in other organisms, data for *B. pertussis* is limited to ZccR, a MerR-like regulator responsive to Zn^2+^, Cd^2+^ and Co^2+^ (Kidd & Brown, 2003). Similarly, Ni^2+^-dependent metalloenzymes are associated with pathogenesis. These include bacterial ureases that neutralize acidic pH to disrupt epithelial barriers, and superoxide dismutases that counteract host oxidative stress (Alfano & Cavazza, 2020; de Reuse, 2017). Therefore, utilizing a high-affinity transport system for Zn^2+^ and Ni^2+^ offers a potential competitive advantage during *B. pertussis* infection.

The uptake of metal-citrate complexes is well established in other transporter families, capitalizing on the abundance of citrate in host tissues and its efficiency as a chelating agent. The Citrate-Metal:Proton Symporter (CitMHS) family uses electrochemical gradients to import citrate chelated to various cations, with members showing selectivity based on ionic radius (Krom *et al*, 2000; Lensbouer & Doyle, 2010; Mortera *et al*, 2013). However, the absence of an associated solute binding protein in this family results in relatively low affinity, ranging in the order of 30 μM to 60 μM. Among SBP-dependent systems, the Fec ABC transporter operon is well characterized for ferric citrate uptake, where the periplasmic protein FecB binds various trivalent cations (Banerjee *et al*, 2016; Mey *et al*, 2021) with the tightest reported citrate-binding affinity at 2.6 nM (Fukushima *et al*, 2012). Other ABC systems, like ZnuABC, transport free Zn²⁺ directly (Cerasi *et al*, 2013). However, the use of ATP for transport incurs a high cost for ion transport. In contrast, the Tripartite Tricarboxylate Transporter (TTT) family balances high-affinity uptake with the energetic efficiency of utilizing electrochemical gradients. Previous studies on BctC homologs highlight how divalent cations influence binding. In *Salmonella typhimurium*, Fe^2+^, Ca^2+^ and Mn^2+^ enhanced TctC binding to radiolabeled citrate by 43%, 26%, and 26%, respectively, whereas affinity decreased with Mg²⁺, Ni²⁺, Zn²⁺, and Co²⁺ (Sweet *et al*, 1979). Our DSF results confirmed that _se_TctC displays higher thermal stabilization in the presence of citrate and Mn^2+^. Similarly, (Brocker *et al*, 2009) reported that citrate uptake in *Corynebacterium glutamicum* increased in the presence of Ca²⁺ and Mg²⁺, though our DSF screening only showed enhanced thermal stability with Mn^2+^. Because DSF is an initial screening tool, a lack of thermal stabilization does not definitively rule out binding, and comprehensive binding assays, such as the ITC data presented here for BctC, are required to establish true affinities. Collectively, these data underscore the widespread adaptation of the high-affinity TTT family to utilize citrate for divalent cation acquisition.

Although citrate is the canonical ligand for Tripartite Tricarboxylate Transporter (TTT) systems, structural determinants of its binding have remained elusive. We determined the BctC crystal structure in three states: apo, open, and citrate/zinc-bound (closed conformation). In the open state, citrate displays partial occupancy and limited electron density, in the vicinity of Phe48 and pincer loop 1 (P41–D49), both in domain 1, while in the closed state, we unambiguously resolved the citrate–Zn²⁺ complex within the binding pocket, coordinated by three water molecules. The transition from open to closed conformation involves a 33.5° rigid-body motion of domain 2, coupled with a subtle reorganization of pincer loop 2 (Phe194–Gly196), mediated by the presence of Zn^2+^. This structural reorganization validates the substrate recognition mechanism proposed by Winnen *et al*. (2003), wherein domain 1 and pincer loop 1 facilitate initial ligand capture, followed by domain 2 closure upon ligand validation. While TTT substrate-binding proteins (SBPs) typically utilize two structural water molecules to bridge the ligand to both pincer loops (Rosa *et al*, 2018), BctC uniquely employs only one (Water 56) to bridge the ligand and structure pincer loop 1. This displacement of the citrate carboxylate group, required to accommodate zinc coordination, prevents the second conserved water molecule (Water 238) from interacting with the ligand, restricting its role instead to stabilizing the inter-domain interface.

The translocation mechanism and transmembrane architecture of the TTT family remain poorly defined. Leveraging *in silico* modeling, we propose that BctCBA operates via an elevator-type symport mechanism, a model likely applicable across the TTT family. Our structural predictions reveal a characteristic topological pseudosymmetry, with distinct elevator and scaffold domains reminiscent of DASS/SLC13 and TRAP transporters (Davies *et al*, 2024; Mulligan *et al*, 2016).

Within the BctA elevator domain, a deeply buried acidic pocket—defined by E75 and Y71 on helix 4 and residues at the tips of mirrored hairpins—forms the putative citrate-binding site. These hairpins, together with conserved G-Hy₃-∗G-Hy₃-∗G-Hy₂-∗P-G-Hy-G motifs (Winnen *et al*, 2003), form a clamshell architecture for Na⁺ coordination. We propose a cooperative binding model where Na⁺ ions stabilize the hairpin loops and partially neutralize the pocket’s negative charge to facilitate citrate docking, analogous to the Na⁺-dependent stabilization of Hp loops observed in VcINDY (Sauer *et al*, 2022).

BctC is essential for high-affinity substrate capture (Antoine *et al*, 2005). Our models show BctC docked at the scaffold region near an acidic periplasmic loop of BctA. This loop may act as a “scooping-loop” to trigger substrate release (Oldham *et al*, 2013) or modulate the solvent network within the BctC binding pocket. Notably, the closed conformation of BctC is only sterically compatible with _in_BctBA, mirroring the conformational coupling observed between SiaP and SiaQM (Peter *et al*, 2024).

At physiological pH, citrate carries a -3 charge. While TRAP transporters are proposed to co-transport three Na⁺ ions despite only two identified binding sites (Davies *et al*, 2024; Fitzgerald *et al*, 2017), the additional transport of a divalent cation in BctCBA would render the process electrogenic. Consequently, we propose a “translocator with an operator” mechanism (Davies *et al*, 2024): closed BctC interacts with the scaffold domain of _in_BctBA, inducing a conformational towards _out_BctBA that triggers BctC opening. An acidic channel spanning BctC and _out_BctA then facilitates the migration of Na⁺ ions and the citrate–cation complex into the _out_BctA pocket. This coordination drives the elevator domain back to the _in_BctBA conformation, resetting the system upon substrate release.

In conclusion, our biochemical and structural characterization of BctC redefines its role from a simple citrate-binding protein to an orchestrator of citrate-mediated divalent cation capture. By capturing the distinct conformational transitions of BctC—from the flexible apo form to the closed, metal-coordinated state—this work provides a structural framework for understanding substrate recognition within the TTT family. Furthermore, our *in silico* modeling offers a plausible structural baseline for the BctCBA transport mechanism, hinting at a potentially electrogenic, elevator-type symport. While the precise dynamics of transmembrane translocation and the *in vivo* implications for *B. pertussis* virulence remain to be experimentally validated, these findings provide critical insights into how pathogens may leverage the TTT family to navigate host nutritional immunity.

## Methodology

### Strains and Molecular biology

*bctC* gene sequence was retrieved from the genome of *Bordetella pertussis* Tohama I (RefSeq assembly accession GCF_000195715.1 gene BP3867). The N-terminal signal was estimated using the SignalP server and removed. The gene was codon optimized and synthesized in pUC57 by GeneScript. *bctC* gene was subclonned into the pNIC-Bsa4 vector via Ligase Independent Cloning, generating a construct from residue 31, with a N-terminal His₆ tag, cleavable by TEV Protease. Primers used were ON_LT47 (forward; 5’-<u>TACTTCCAATCCATG</u>GCGGCCGGCGACGAG-3’) and ON_LT40 (reverse; 5’-<u>TATCCACCTTTACTG</u>TTACTTCTTGATGAGGCCGAAGCTGTCG-3’), and transformed into *E. coli* DH5α cells.

BctC homologs sequences followed the same procedure, except that they were synthetized by FastBio directly into pET28a, with an N-terminal His₆ tag cleavable by TEV Protease. Uniprot entries used as template are: _SE_TctC: Q9FA46, residues 23-324; _CG_TctC: Q8NLW1 residues 43-334; _GT_TctC: A4IPE9 residues 22-342 ; _AV_TctC: W0PK92 residues 29-331; _PA_TctC: A0A0H2Z6Z0 Residues 26-327. NCBI accession codes for optimized codons are: BctC: PZ631086; _SE_TctC: PZ631087; _CG_TctC: PZ631088; _GT_TctC: PZ631089; _AV_TctC:PZ631090; _PA_TctC: PZ631091.

### Protein overexpression and purification

BctC and TctC homolog encoding constructs were transformed into *Escherichia coli* BL21 (DE3) cells for recombinant protein expression. Cells were grown in LB medium supplemented with 50 µg mL⁻¹ kanamycin at 37 °C with shaking at 180 rpm until reaching an OD₆₀₀ of 0.7. Protein expression was induced by the addition of 0.1 mM IPTG, and cultures were subsequently incubated overnight at 16 °C with shaking at 180 rpm before harvesting by centrifugation. Cell pellets were resuspended in 30 mL of binding buffer (20 mM HEPES, 300 mM NaCl, 20 mM imidazole, pH 7.4) and lysed by sonication (3 s ON/10 s OFF cycles) at 30% amplitude for 5 min while maintained on ice. The lysate was clarified by centrifugation at 13,000 × *g* for 35 min at 4 °C. The resulting cell-free extract was filtered and loaded onto a 5 mL HisTrap™ HP affinity chromatography column (Cytiva) pre-equilibrated with binding buffer. Recombinant BctC was eluted using a linear 0–500 mM imidazole gradient in the elution buffer (20 mM HEPES, 300 mM NaCl, 500 mM imidazole, pH 7.4). To remove any co-purified ligands, the eluted BctC protein was subjected to a denaturation–refolding procedure. The protein was first dialyzed against 20 mM HEPES, 300 mM NaCl, pH 7.4, supplemented with 6 M urea, using SnakeSkin™ dialysis tubing under gentle agitation for 12–16 h at 4 °C. Refolding was then achieved through a stepwise decrease in urea concentration from 6 M to 0 M in the same buffer, with dialysis solutions replaced every 2 h while maintaining agitation at 4 °C. Following refolding, the N-terminal His-tag was removed by digestion with TEV protease at a 1:6 (protease:protein, w/w) ratio during overnight dialysis in HEPES-NaCl buffer (pH 7.4) at 4 °C. The sample was subsequently passed through a gravity-flow column containing Ni²⁺-charged affinity resin pre-equilibrated with the same buffer. The tag-free BctC protein was recovered in the flow-through fraction. Final purification was performed by size-exclusion chromatography (SEC) using a Superdex™ 75 Increase 10/300 GL column (Cytiva) equilibrated with 50 mM HEPES, 150 mM NaCl, and 1% (w/v) glycerol (pH 7.4). Chromatography was carried out at a flow rate of 0.65 mL min⁻¹ using an ÄKTA go system (Cytiva). Protein purity was assessed by SDS-PAGE. Fractions containing BctC were pooled and concentrated using Amicon® Ultra-15 centrifugal filter units with a 10 kDa molecular weight cut-off (Merck Millipore). A calibration curve was determined for the Superdex™ 75 Increase 10/300 GL column (Cytiva) using the Gel Filtration Calibration Kit LMW (low molecular weight) with the following standards: conalbumin (75 kDa), ovalbumin (44 kDa), carbonic anhydrase (29 kDa), Ribonuclease A (13.7 kDa), aprotinin (6.5 kDa), and Blue Dextran 2000, yielding the linear regression equation f(x)=−0.387x+1.971 with an R2=0.995. When required, the buffer was supplemented with ZnSO_4_ 1 mM and sodium citrate 1 mM.

### Circular Dichroism of BctC

Prior to analysis, the protein sample was centrifuged at 13,000 rpm for 10 min, and the concentration was measured and adjusted before CD measurements. Proteins were diluted in urea-free buffer (20 mM HEPES, 300 mM NaCl, pH 7.4) to a final concentration of 5 μM. CD measurements were performed on a J-720 spectropolarimeter (JASCO Corporation), and spectra were recorded in the far-UV region from 260 to 200 nm using a 1 mm pathlength quartz cuvette, ensuring detector voltages remained below 700 V and using a detector sensitivity of 100 mdeg. The obtained spectra were compared with characteristic secondary structure profiles, including α-helices, β-sheets, and disordered structures, to confirm recovery of the native protein conformation.

### Differential Scanning Fluorimetry

Thermal stability of BctC was assessed by differential scanning fluorimetry against a ligand library composed of 85 compounds (Appendix Table 1). The assays were performed in 384-well plates, with each reaction containing 2 μM protein, 5x SYPRO Orange dye (Thermo Fisher Scientific), and 50 µM of ligand in HEPES-NaCl buffer were heated up to 95 °C using a QuantStudio^TM^ 6 Flex Real-Time PCR (Applied Biosystems) system. Melting temperatures were determined from the first derivative of the fluorescence curves and ligands that exhibited a ΔTm greater than 2 °C were subjected to further affinity studies. To investigate the effect of metal ions on ligand binding, additional DSF assays were performed using citrate alone, divalent cations (Ca^2+^, Fe^2+^, Zn^2+^, Mg^2+^, Mn^2+^ and Ni^2+^) and citrate in combination with each cation. Citrate and metal ions were tested at a final concentration of 50 µM, with pH adjusted prior to analysis. These experiments were conducted under the same conditions used for the ligand library screening. Statistical analyses were performed using Welch’s t-test (two-tailed), assuming unequal variances between groups. Each cation condition was compared with the buffer control, and each citrate + cation conditions were compared with citrate control. Differences were considered statistically significant when p < 0.05.

### Isothermal Titration Calorimetry

ITC experiments were performed at 25 °C using a MicroCal PEAQ-ITC Automated (Malvern Panalytical) calorimeter, available at the National Laboratory of Bio-renewables (CNPEM, Brazil). Protein samples were supplemented with 5mM divalent cation and dialyzed against HEPES-NaCl buffer (pH 7.4), subsequently adjusted to a final concentration of 120 µM. The sample cell was loaded with the protein and cation solution, while the injection syringe was filled with 600 µM citrate prepared in the same dialysis buffer. Titrations consisted of an initial 0.4 µL injection followed by 19 injections of 2 µL each, with 150 s intervals between injections and a stirring speed of 750 rpm. Control experiments were performed by titrating the protein buffer under identical experimental conditions. The resulting thermograms were analyzed using MicroCal PEAQ-ITC Analysis Software (Malvern Panalytical) assuming a one-site binding model.

### Crystallography and X-ray diffraction

Protein was concentrated to 15 mg mL⁻¹ (to test its holo state, 1 mM ligands were added). A series of crystallization conditions were tested using commercial crystallization screening kits (Molecular Dimensions), in 96-well plates. The sitting-drop vapor diffusion method was used, with 100 nL of protein and 100 nL of precipitating solution per drop, dispensed by automated pipetting using Mosquito® equipment (SPT Labtech) and the plates were incubated at 19 °C.

Diffraction data of BctC crystals were collected at Manacá beamline at Sirius (LNLS, Brazil) and processed using autoPROC (Vonrhein *et al*, 2011). The structure was solved by molecular replacement and refined using standard crystallographic software (Agirre *et al*, 2023; Krissinel *et al*, 2018; Liebschner *et al*, 2019), Phaser (McCoy *et al*, 2007) and Coot (Emsley *et al*, 2010). Data collection and refinement statistics are summarized in Table 2. Metal coordination analysis were performed using the CheckMyMetal server (Gucwa *et al*, 2023). Structure analysis and figure generation were performed in ChimeraX (Pettersen *et al*, 2004).

### *In Silico* analysis of TctCBA

*In silico* modelling of BctCBA was done using the Alphafold3 server (Abramson *et al*, 2024), with default parameters, and one copy of each protein, one citrate and two sodium ions as input. Structure analysis and figure generation were performed in ChimeraX (Pettersen *et al*, 2004).

## Data Availability

BctC crystallographic structures are available in the Protein Data Bank, under accession codes 9ZPR (Apo 16 ASU), 37CE (Apo), 37EH (Open), 37KY (closed, bound to citrate and zinc).

Codon-optimized sequences for BctC and TctC homologs are available in the National Center for Biotechnological Information, under the following accession codes: BctC: PZ631086; _SE_TctC: PZ631087; _CG_TctC: PZ631088; _GT_TctC: PZ631089; _AV_TctC:PZ631090; _PA_TctC: PZ631091.

## Authors contributions

Experimental Design - RJB, MVM, LTR

Molecular Biology - LBC, MSD, LRS, KBM, MHB

Protein expression and purification - LBC, MSD, LRS, NFB, KBM, MHB, LTR

DSF analysis - LBC, MSD, LRS, KBM, LTR

ITC analysis - LBC, MSD, LTR

Protein crystallography - LBC, MSD, JPA, RJB, MHB, LTR

In silico model analysis - NFB, GGS, LTR

Manuscript writing - LBC, LRS, MVM, RJB, LTR

## Acknowledgements

We thank Carlos Henrique Inacio Ramos and Natália Galdi Quel for their support with circular dichroism analysis.

This study was financed by the São Paulo Research Foundation (FAPESP), Brasil. Process Numbers 2022/11936-6 and 2023/05408-0 to LTR, 2021/10577-0 to MVM , GGS and LTR, 2024/09522-4 to LBC and 2024/20003-9 to NFB.

This study was financed in part by the Coordenação de Aperfeiçoamento de Pessoal de Nível Superior - Brasil (CAPES) - Finance Code 001. 88887.103984/2025-00.

This research used facilities of the Brazilian Synchrotron Light Laboratory (LNLS), Brazilian Biorenewables National Laboratory (LNBR) and Brazilian Biosciences National Laboratory (LNBio), all part of the Brazilian Center for Research in Energy and Materials (CNPEM), a private non-profit organization under the supervision of the Brazilian Ministry for Science, Technology, and Innovations (MCTI). The LNLS Manacá beamline staff is acknowledged for the assistance during the experiments 20241239 and 20253736. The LNBR BFM-ITC staff is acknowledged for the assistance during the experiments 20242938. The LNBio Robolab staff is acknowledged for the assistance during the experiments 20254249. The AI tools ChatGPT and Gemini Academy were used for English grammar improvement only.

## Figure Legends

**Figure Expanded View 1**. Stability characterization of BctC. (a) Differential scanning fluorimetry (DSF) analysis of BctC across a range of pH conditions. (b) Circular dichroism spectra of BctC before urea-induced denaturation (black) and after refolding by renaturation (red).

**Figure Expanded View 2.** Isothermal titration calorimetry analysis of citrate binding to BctC in the presence of divalent cations. Representative isothermal titration calorimetry (ITC) profiles of citrate titrated into BctC in the presence of different divalent cations, illustrating the effects of metal ions on ligand binding.

**Figure Expanded View 3**. Structural basis of citrate recognition and binding-site organization in BctC. (a) Three-dimensional view of BctC open structure binding cavity. Oxygen atoms are shown in red, nitrogen atoms in blue, water molecules as red spheres. A citrate molecule was inserted in the figure for reference, with carbon atoms colored gray. Carbon atoms from BctC residues belonging to Domain 1 are shown in olive green, whereas those from Domain 2 are shown in orange. The polder map is displayed contoured at 4.5 RMSD. (b) BctC closed structure, colored as in Fig. 2B. The α-helices surrounding the binding pocket are shown in a rainbow gradient ranging from the N-terminus (blue) to the C-terminus (red), highlighting the convergent orientation of their N-terminal ends toward the ligand-coordination site.

**Figure Expanded View 4.** In silico structural models of the BctCBA complex and analysis of substrate-transfer interfaces. (a) In silico models of the BctCBA complex colored according to pLDDT confidence scores, together with their corresponding Predicted Aligned Error (PAE) plots. _in_BctCBA (left) depicts BctA in a resting, cytoplasm-facing conformation, whereas _out_BctCBA (right) depicts BctA in a periplasm-facing conformation. (b) Surface representation of the BctA elevator domain, colored according to electrostatic potential, with acidic regions shown in red, basic regions in blue, and neutral regions in white. A prominent acidic pocket is observed at the center of the domain and is predicted to correspond to the citrate-binding site. (c) Alignment of the closed conformation of BctC onto _out_BctCBA. The original open-state BctC from the model is omitted for clarity. The closed-state BctC is shown as a white surface outline, with BctA regions located within 2 Å of BctC highlighted in red. Alignments were performed using either Domain 1 (left) or Domain 2 (right) of the modelled open BctC structure as the reference.

## Table Legends

**Appendix Table 1:** Components of the Differential Scanning Fluorescence screening library.

**Appendix Table 2:** Individual thermodynamic parameters obtained from independent ITC experiments.

